# Distinct genomic adaptations of the methanogenic archaeal genus *Methanocorpusculum* to symbiosis with animals and protists

**DOI:** 10.64898/2026.05.13.724943

**Authors:** Anna Schrecengost, Johana Rotterová, Ivan Čepička, Roxanne A. Beinart

## Abstract

Most of our understanding of endosymbiosis originates from the bacterial endosymbionts of multicellular, terrestrial hosts, which represent habitats with dramatically different selective pressures than inside a protist cell. Methanogenic archaea from the genus *Methanocorpusculum* are among the few known intracellular archaea and form unique symbioses with both animal and protist hosts, providing a unique opportunity to contrast symbiont evolution and function in very distinct host types. Here, we conducted phylo- and pangenomic analyses on 106 *Methanocorpusculum* strains originating from animal and ciliate hosts as well as environmental habitats. We recovered two divergent clades corresponding to animal gut-associated and intracellular ciliate-associated/environmental lineages and found that ciliate-associated and environmental *Methanocorpusculum* are virtually indistinguishable functionally and phylogenetically. Ciliate-associated symbionts retained broad biosynthetic capacity and encoded functions related to osmotic stress tolerance and adhesion within the host cell, while animal gut-associated symbionts exhibited patterns of genome streamlining and nutrient scavenging consistent with host supply and immune adaptation. Our findings illuminate how the contrasting selective pressures of protists and animal hosts have driven divergent evolutionary and functional strategies in congeneric archaeal symbionts.

## Introduction

Microbial symbiosis amongst prokaryotes and eukaryotes was a driving force in the evolution of complex life, from the first eukaryotic cell to the development of multicellular organisms (1,2). Today we understand that microbial symbioses profoundly shape the ecology and evolution of both partners, as well as generate emergent properties and functions (3,4); however, the majority of this understanding comes from the bacterial symbionts of terrestrial, multicellular hosts, leaving archaeal endosymbionts and protist symbioses relatively unexplored (5–7). Methanogenic archaea are widespread, diverse, and ecologically significant symbionts: virtually all vertebrates and a diversity of arthropods host methanogens in their digestive tracts, contributing around 20% of total annual methane emissions (6,8–11), and the methanogenic endosymbionts of anaerobic protists represent some of the only known intracellular archaea (6,12). Despite this, virtually nothing is known about how the distinct selective pressures of their varied host environments shape their evolution and function.

To better understand the ecology and evolution of microbial symbiosis across eukaryotic diversity, we can compare microbes associated intracellularly with unicellular protists and extracellularly with multicellular eukaryotes. Establishing symbiosis is known to be a complex process in multicellular hosts, requiring colonization of the host environment, competition with other microbes, immune evasion, and transmission to the next generation (13). While symbionts presumably face comparable challenges in protists, the mechanisms enabling prokaryotes to reside within unicellular hosts are understudied (14).

*Methanocorpusculum* is a genus of methanogenic archaea that is harbored extracellularly within animal gastrointestinal tracts and intracellularly inside anaerobic marine and freshwater ciliates (16–21), as well as in habitats such as wastewater and anaerobic digesters (22). The methanogenic endosymbionts of anaerobic protists are among the only known intracellular archaea, representing an excellent opportunity to better understand the evolutionary and functional dynamics of endosymbiosis (12). They were likely acquired during host transitions to anaerobiosis, facilitating host’s permanent residence in hypoxia via the syntrophic consumption of host metabolic waste products (e.g., H_2_) during symbiont methanogenesis (23–25). These symbionts are both host- and habitat-specific, exhibiting stability at the host species level via vertical transmission, but there is evidence of symbiont replacements at higher taxonomic levels (19–21,26).

Within the gastrointestinal tracts of animals, methanogenic archaea are ubiquitous and diverse. Similar to those in protists, they balance H_2_ concentrations, promoting the growth of co-occurring fermentative bacteria and increasing host energy yield: in ruminants, fermentation products provide ∼70% of host energy (27). Animal-associated microbes and methanogens often show evidence of co-evolution with their hosts (28–30). Gut colonization occurs very soon after birth, largely through vertical transmission in vertebrates, although in invertebrates horizontal acquisition from the environment may play a larger role (8,9,27). Establishment and maintenance of gut-associated methanogens is influenced by a combination of host genetics, diet, immune status, and selective pressure on the host (27,31). Gut-associated methanogens themselves can contribute to host specificity through adaptations such as glycan mimicry and host attachment (32,33).

Therefore, this genus of methanogens, which forms unique, mutualistic symbioses across the eukaryotic tree of life, provides an opportunity to investigate how distinct host associations shape symbiont ecology and evolution. Interestingly, *Methanocorpusculum* is one of few prokaryotic genera that form mutualistic associations with both metazoans and protists. Most other examples of symbionts harbored by both animals and protists involve pathogenic prokaryotes and/or endobiotic protist hosts (15,34,35). To our knowledge, methanogens such as *Methanocorpusculum* and *Methanobacterium* are among the only known prokaryotes that occur as both extracellular animal symbionts and intracellular symbionts of free-living protists (6,16).

In this way, *Methanocorpusculum* offers a unique opportunity to examine how symbionts evolve under different selective regimes and transmission strategies. Here, we conducted phylo-and pangenomic analyses on new and existing *Methanocorpusculum* genomes, comparing functional and phylogenetic patterns of symbionts across ciliate hosts, animal guts, and environmental isolates. This effort has revealed how environmental conditions, host phylogeny, and mixed-mode transmission shape the evolution and function of symbiotic *Methanocorpusculum* across disparate host and environmental habitats.

## Experimental Procedures

### Single ciliate cell isolation and DNA sequencing

Single ciliate cells from *Metopus* strains and one *Plagiopyla* strain (JUMA2P) were processed previously (21). Here, we isolated six additional *Plagiopyla* strains (Table S1) using the same protocol. Briefly, cells were isolated by micropipette, washed, and subjected to whole genome amplification (RepliG Advance DNA Single Cell kit, Qiagen). Amplified DNA was quantified (Qubit) and sequenced on Illumina NovaSeq6000 S4 (2×150 bp, SeqWell plexWell™ 96 NGS library kit)

### Assembly and binning of endosymbiont MAGs and dataset curation

Illumina paired-end reads were trimmed with and quality filtered Trimmomatic (v0.39) (36), contaminant reads removed with Bowtie2 (v2.5.2) (37), and paired end reads assembled *de novo* using metaSPAdes (v3.15.5) (38) Contigs were binned with metaBAT2 (v2.15) (39) and MaxBin2 (v2.2.7) (40), and the resulting bins were refined using DAS Tool (v1.1.7) (41), producing consensus metagenome-assembled genomes (MAGs). Full details can be found in the Supplemental Information.

Symbiont MAGs were identified using GTDB-tk classify_wf (42). Based on previous knowledge of these symbiont populations, bins assigned to *Methanocorpusculum* were considered symbiont bins. Only MAGs with ≥80% completeness and ≤5% contamination, as determined by CheckM2 (43), were retained (Table S2). Additional ciliate-associated MAGs were retrieved from Rotterova et al. (21) and Lind et al. (44). Additional environmental and animal gut-associated reference genomes and high-quality MAGs were retrieved from GenBank based on previous phylogenomic analyses of *Methanocorpusculum* (16,17). We excluded a clade reclassified as *Methanorbis* by Protasov et al. in 2023 (17).

### Pangenomic analyses of Methanocorpusculum symbionts and of the family Methanocorpusculaceae

Pangenomic analyses were conducted with anvi’o 8 (45,46), including the generation of contigs databases with anvi-gen-contigs-database and identification of single-core copy genes (SCCGs) and ribosomal markers with anvi-run-hmms. Functional annotation was performed using anvi-run-ncbi-cogs (COG database) and anvi-run-kegg-kofams (KEGG orthologs). KEGG module completeness was estimated using anvi-estimate-metabolism (47). Functional enrichment across host types/lifestyle was computed using anvi-compute-functional-enrichment-in-pan (48), which calculates enrichment scores and associated p- and q-values. We considered any function/module with a q-value <0.05 enriched, and those with enrichment scores >30 highly enriched. Pairwise ANI was calculated using anvi-compute-genome-similarity using the pyani ANIb method.

To better understand the evolutionary context in which functional shifts between animal-and protist-associated genomes occurred, we constructed an additional pangenome including Methanocorpusculaceae (*Methanocorpusculum, Methanorbis,* and *Methanocalculus*) and representatives from its most closely-related genera, *Methanofollis* and *Methanoculleus* (17,22). One high-quality representative ciliate-associated MAG per host strain was included alongside environmental and animal gut-associated reference genomes/MAGs (Table S5). Pangenomic analysis was conducted with anvi’o 8 as described above, except we used –minbit 0.3 rather than the default 0.5.

### Phylogenomic analyses of symbionts and phylogenetic analysis of hosts

We inferred a maximum-likelihood phylogenomic tree based on the concatenated alignment of 220 single-copy core genes (SCCGs) identified across the pangenome of *Methanocorpusculum* MAGs/genomes using IQ-TREE with 1000 bootstraps under the WAG model. A second tree including Methanomicrobiales was constructed similarly (199 SCCGs across 86 genomes/MAGs).

Host ciliate phylogenies were inferred from previously obtained 18S rRNA gene sequences (19,20), and additional sequences were recovered via BLASTn from metagenomic assemblies (BLAST+ v2.15.0) (49). Sequences were aligned using the MAFFT algorithm (G-INS-i), manually trimmed to the primer regions using AliView (50), and phylogenetic trees generated using a ML method in RAxML (GTRGAMMAI, 1000 bootstraps (51)).

## Results

### Genome-resolved diversity and phylogenomics of Methanocorpusculum

Here, we present new ciliate-associated *Methanocorpusculum* MAGs which were generated from single-cell metagenomic sequencing of 25 total cells isolated from five *Plagiopyla* strains (Table S1), resulting in one high-quality *Methanocorpusculum* symbiont genome per cell. The MAGs were 79.84-94.12% complete (average 89% ±4.32% s.d.), with contamination 0.42% (±0.432% s.d.) and GC content between 0.44-0.5 (Table S2). Genome sizes ranged between 1.35 Mb - 2.28 Mb and contained 1533-1994 protein-coding genes. Together with the *Plagiopyla* strain from (21), this data represent endosymbiont genomes from six host species-level lineages (19) (Figure 1). We also include previously described *Methanocorpusculum* endosymbiont genomes from ten *Metopus* strains, representing two host species (21) (Figure 1). In total, we included 106 *Methanocorpusculum* genomes/MAGs in our analysis: 84 ciliate-associated MAGs (34 *Plagiopyla*, 50 *Metopus*), nine environmental, and 13 animal-associated (Table S2).

**Figure 1.**
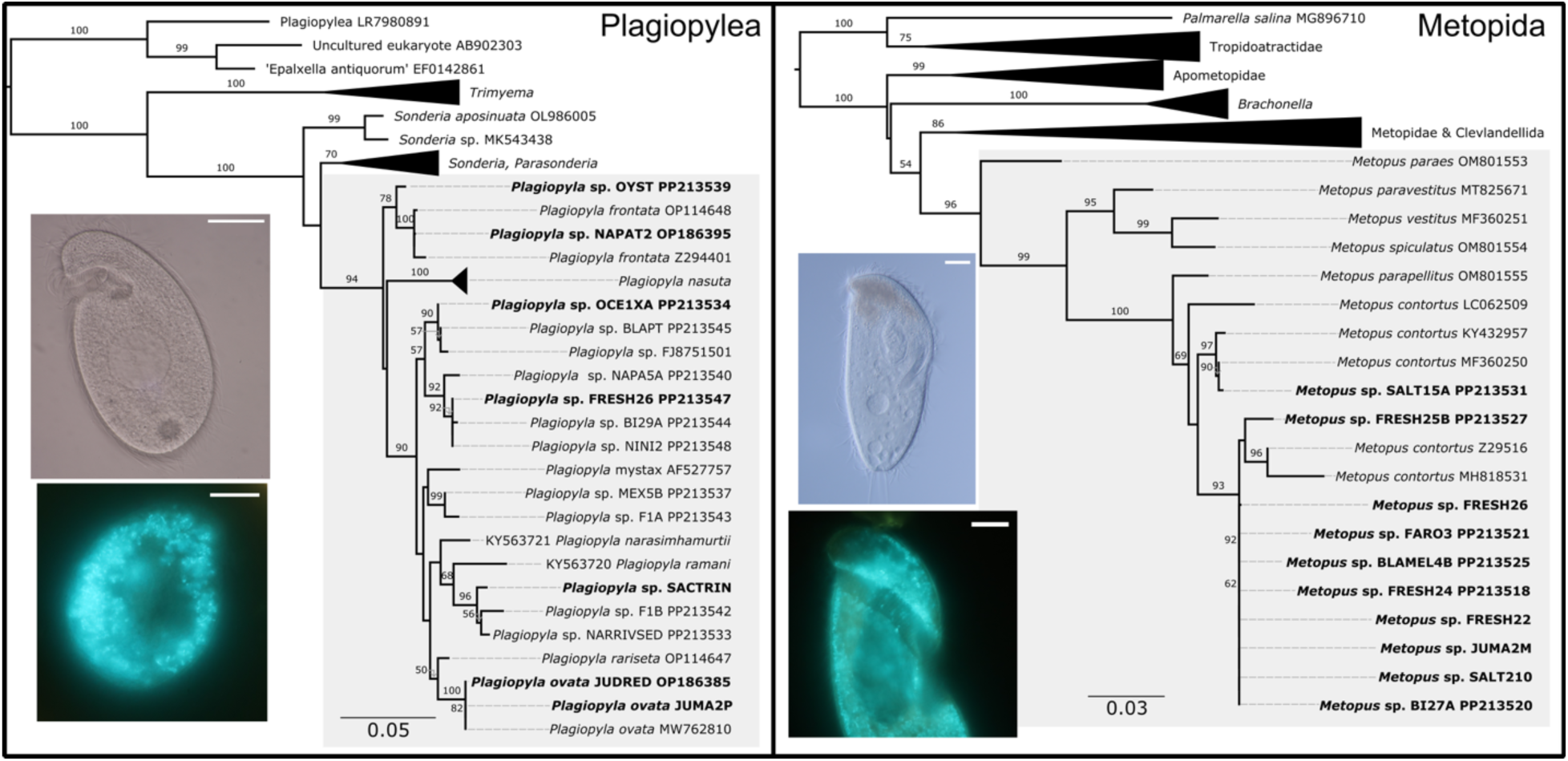
Phylogenetic trees and micrographs of *Plagiopyla* ciliates (left) and *Metopus* ciliates (right) whose symbionts were investigated in this study. Trees constructed from 18S rRNA gene sequences of the strains in this study (shown in bold), as well as published sequences from cultured representatives and a few environmental sequences from the Class Plagiopylea and the Order Metopida. Representative light micrographs of *Plagiopyla* shown at left: *Plagiopyla frontata* NAPAT2 shown on top, *Plagiopyla* sp. OYST shown at bottom with blue-green autofluorescence of methanogenic endosymbionts. Light micrographs of representative *Metopus* shown at right: *Metopus* sp. FRESH25B shown on top, and *Metopus* sp. FRESH24 shown at bottom. Scale bar = 20 μm.

Our phylogenomic analysis recovered two well-supported *Methanocorpusculum* clades: one containing animal-associated and the other containing ciliate-associated and environmental lineages. Symbionts from four *Plagiopyla* species (*P. ovata* JUDRED, *P. frontata* NAPAT2, OYST, and JUMA2P) formed a well-supported clade, as did all *Metopus*-associated symbionts plus three *Plagiopyla*-associated symbionts (SACTRIN, F26, OCE1XA) (Figure 2A). Most environmental genomes formed a well-supported, long-branching clade in our phylogeny, with a few interspersed among ciliate-associated lineages (*M.* sp. GCF30655665 and *M. labreanum* Z). The animal-associated and ciliate-associated/environmental clades were also genomically distinct, sharing 72.89% similarity (±0.004 s.d.) based on the ANI of all alignable regions in the genome, which is a low value for cogeners but consistent with assignment to *Methanocorpusculum* based on the analysis of Protasov et al. (2023) and on taxonomic assignment with GTBD-tk (Table S2).

**Figure 2.**
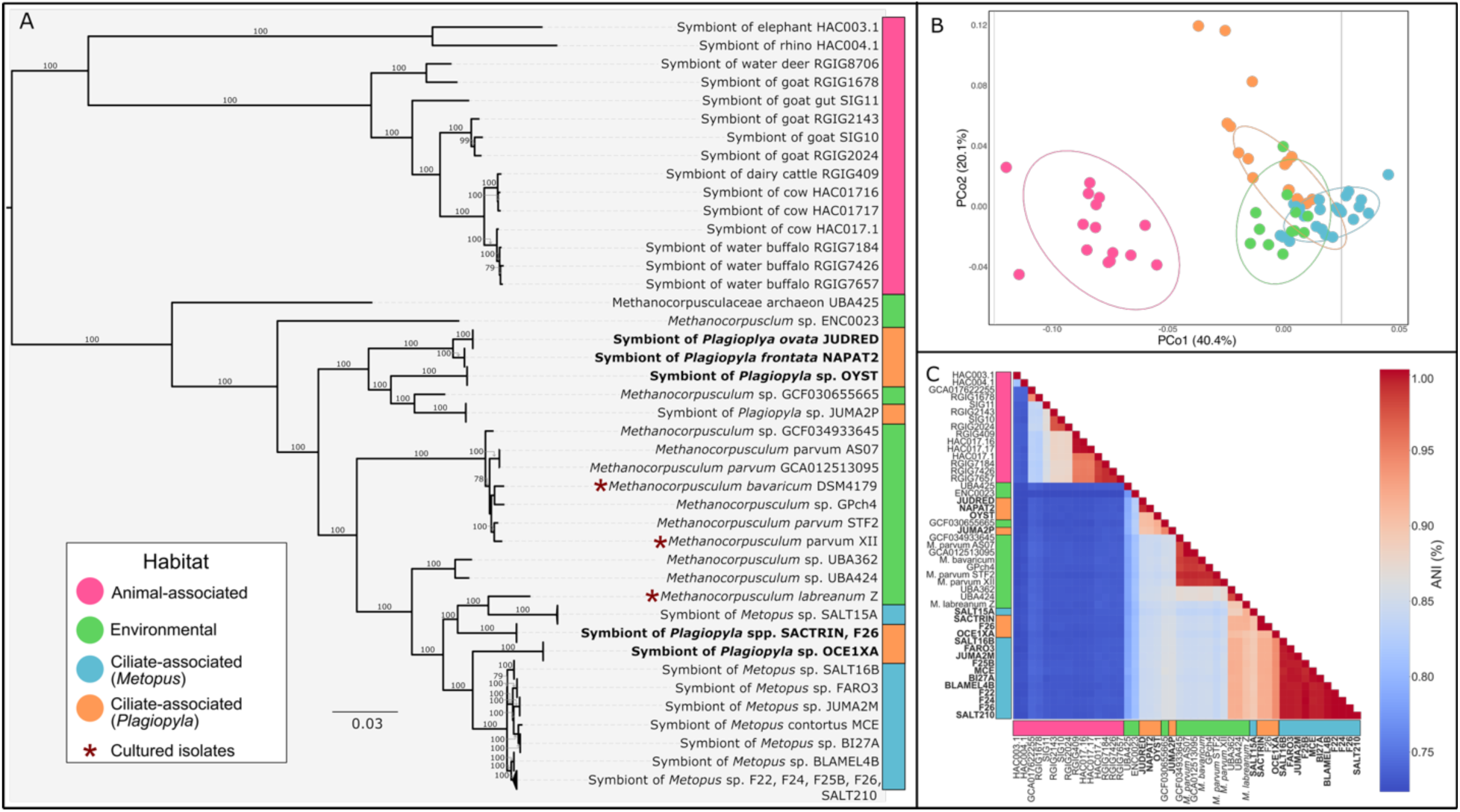
Summary of relationships between *Methanocorpusculum* from different habitats. Colors in the legend indicate host habitat for symbiotic species (ciliate-associated, “*Metopus”* and “*Plagiopyla”*, and “Animal-associated”) and free-living species are labeled “Environmental”. (A): A maximum-likelihood phylogenomic tree was inferred from a concatenated alignment of 220 single-copy core genes (SCCGs) identified across the *Methanocorpusclum* pangenome and constructed using 1000 bootstraps (only bootstrap values above 70 are shown on branches). Scale bar indicates substitutions per site. Colored bars alongside node labels indicate habitat, as pictured in the legend. Genomes originating from cultured populations are labelled with red asterisks, and strains from this study are in bold. (B): PCoA analysis based on Bray-Curtis distances calculated from pathway completeness profiles of KEGG modules in the pangenomes. Points represent genomes or MAGs and are colored by habitat. Axes show percentage of variance explained: PCo1: 40.4%, PCo2: 20.1%. Ellipses represent 95% confidence interval for grouping by habitat. (C): Heatmap illustrating average nucleotide identity (ANI) between genomes/MAGs. Color scale at right indicates ANI percentage of pairs. Colored bars alongside the x- and y-axis indicate habitat of *Methanocorpusculum* populations.

Based on ANI (Figure 1C, Table S3), taxonomic assignment with GTDB-tk (Table S2), as well as our phylogenomic analysis (Figure 1A), the ciliate-associated MAGs comprise six *Methanocorpusculum* species across two host genera: four associated with six *Plagiopyla* species and two with two *Metopus* species.

### Methanocorpusculum metabolic potential across diverse host-associated and free-living lifestyles

We conducted a pangenomic analysis of 106 *Methanocorpusculum* genomes/MAGs to assess functional differences across lifestyles and host types. The pangenome contains 192,845 genes across 6,881 gene clusters, with a core genome (present in 100% of the genomes) of 540 genes (7.8%), an accessory genome of 3406 (49.5%), and 2935 singletons (genes only present in a single MAG, 43%). The core genome is larger than a previous analysis of the pangenome of animal-associated *Methanocorpusculum* and *Methanorbis* (189 genes, 1.8%) (16).

Ordinations of KEGG module completeness (Figure 2b) revealed that the symbiont genomes clustered based on host type: animal-associated lineages were distinct, while ciliate-associated and environmental genomes clustered together. Subsequent functional enrichment analyses (COG and KEGG; Table S3; Figures 3-4) revealed that the metabolic divergence observed across host type was likely driven differences in enzymes/pathways related to the biosynthesis of essential amino acids and cofactors, as well as nutrient scavenging (Table S3, Figures 3 and 4). Ciliate-associated/environmental genomes were highly enriched in genes involved in amino acid and cofactor biosynthesis (e.g. tryptophan: *trpA*, *trpB*, *trpE*, *trpD*; leucine: *leuA, leuB leuC, leuD*; methionine: *hcybio, met2, met17*; KEGG modules for phenylalanine, tyrosine, leucine, isoleucine, and methionine) as well as modules related to cofactor biosynthesis, including NAD, pyridoxal-P (vitamin B6), heme, and coenzyme F430 biosynthesis. In contrast, animal-associated *Methanocorpusculum* were enriched in several nutrient-salvage and catabolic gene families that were absent (e.g. histidine transporter *hisJ* and *hisM*; di-/tri-peptidase gene *pepD2*; serine/threonine transporter gene *sstT*) or largely absent (e.g. nucleoside transporters *upp, yhhQ, hptA*, the vitamin B6 salvage gene *pdxK*, threonine aldolase *gly1*, and phenylpyruvate tautomerase *pptA*) from other groups.

**Figure 3.**
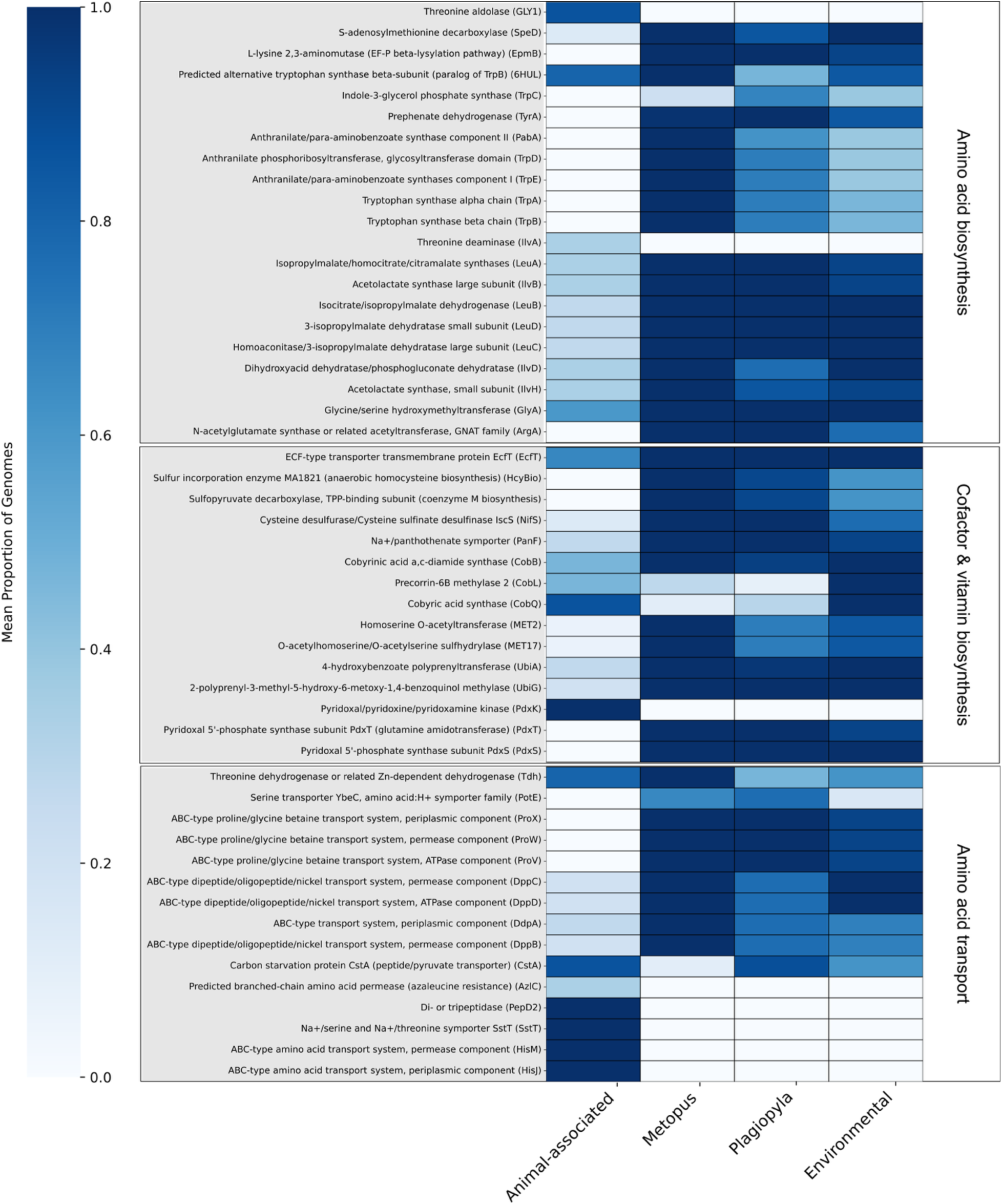
Functionally enriched enzymes involved in amino acid metabolism across host-associated and environmental habitats. Gene clusters were annotated with COG functions. Color indicates the proportion of genomes in each group which contain a given COG function. Only enzymes with a functional enrichment score of at least 30 were included, and all of these are significantly enriched (*p*<0.05).

**Figure 4.**
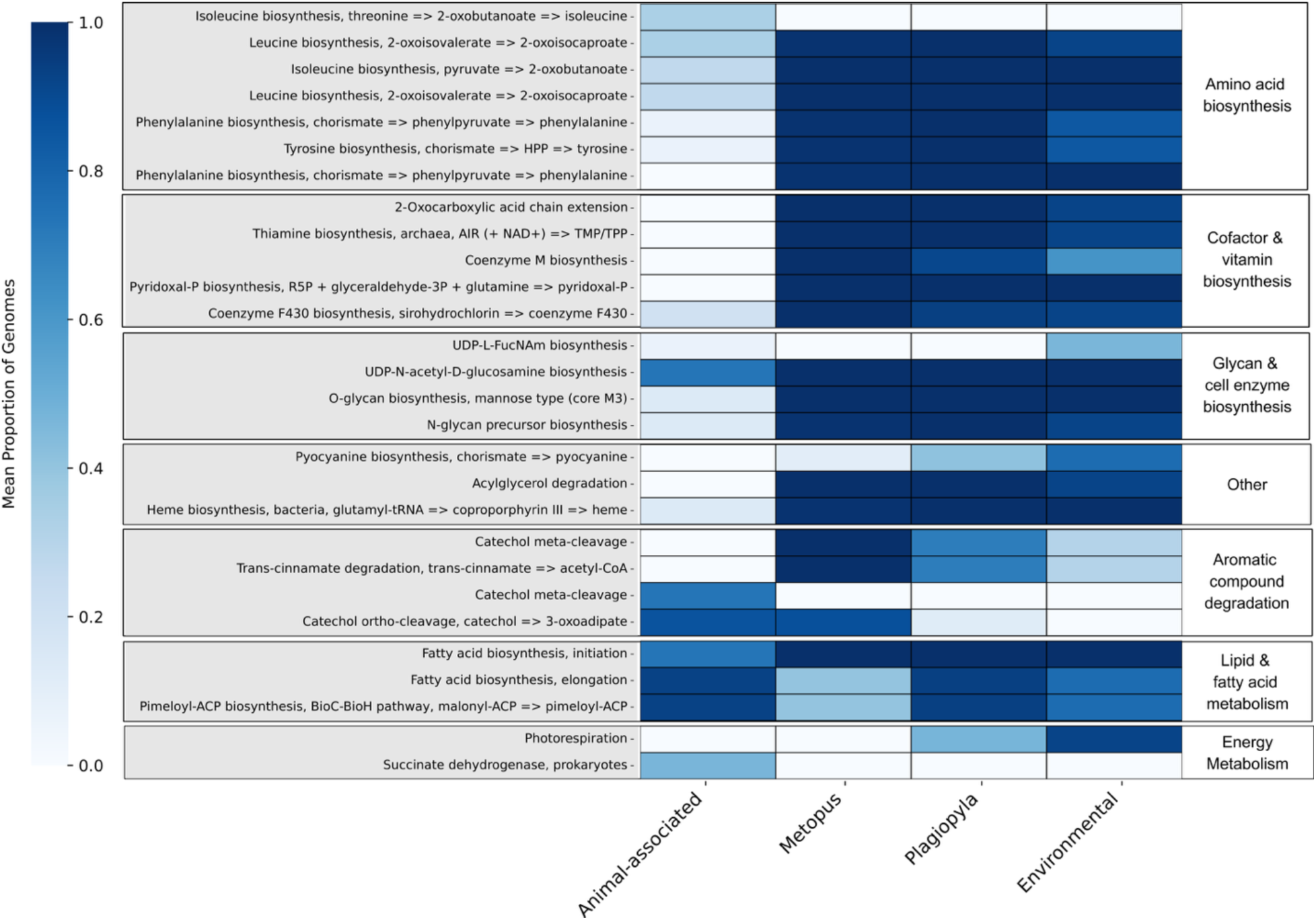
Functionally enriched KEGG modules across host-associated and environmental habitats. Color indicates the proportion of genomes in each group which contain a given KEGG module. Only modules with a functional enrichment score of at least 30 were included, and all of these are significantly enriched (*p*<0.05).

Additionally, ciliate-associated/environmental genomes were fully enriched in genes for proline/glycine transport (*proX, proV, proW*; Figure 3) and highly enriched in KEGG modules associated with glycan biosynthesis (O-glycan biosynthesis, mannose type and N-glycan precursor biosynthesis, Figure 4). Animal-associated genomes uniquely encoded functions related to bile acid detoxification and stress response (3alpha-hydroxycholanate dehydrogenase, the ssDNA/RNA exonuclease *tatD*).

We extended this analysis to also include genomes from closely-related, free-living methanogen lineages (Figure S1). Specifically, we constructed an additional pangenome and phylogenomic tree which included all of Methanocorpusculaceae (*Methanocorpusculum, Methanorbis* (17), and *Methanocalculus*) and its most closely-related genera, *Methanofollis* and *Methanoculleus* (17,22) (Figure S1). In this analysis, ordinations of KEGG module completeness (Figure S2) revealed three major metabolic clusters: free-living *Methanofollis, Methanoculleus,* and *Methanocalculus*; *Methanorbis* and ciliate-associated and environmental *Methanocorpusculum*; and animal-associated *Methanocorpusculum*.

Subsequent functional enrichment analyses of COG functions (Figure S3) and KEGG modules (Figure S4) showed similar results to the *Methanocorpusculum*-only analysis: genes involved in the biosynthesis of tryptophan and osmotic stress tolerance (*proX* and *proW*) were absent in animal-associated genomes but present in the rest of the groups (Figure S3, Table S4). Similarly, pepD2 and the transporter genes *hisM*, *hisJ*, *hptA* were enriched in these animal-associated genomes. On the other hand, there were several functions that were reduced only in the animal-associated *Methanocorpusculum* but present in the rest: biosynthetic leucine and isoleucine genes, *tatD*, *gly1*, valine and isoleucine biosynthesis KEGG modules, and KEGG modules for pyridoxal-p biosynthesis and coenzyme F430 biosynthesis. Additionally, cobalamin biosynthetic KEGG modules and COG enzymes were reduced in animal- and ciliate-associated *Methanocorpusculum*.

## Discussion

### Evolutionary relationships between ciliate-, animal-associated and environmental Methanocorpusculum lineages

Our phylogenomic analysis recovered two divergent, well-supported *Methanocorpusculum* clades: one containing only extracellular, animal-associated lineages and the other comprising intracellular, ciliate-associated *Methanocorpusculum* together with environmental lineages. These two clades are distinct phylogenetically and based on ANI, likely reflecting divergent lifestyles and evolutionary trajectories: animal-associated lineages live extracellularly within their mammalian hosts’ digestive tracts, whereas ciliate-associated species reside intracellularly within anaerobic ciliate cells (30). In contrast to the distinct animal-associated clade, the ciliate-associated lineages form several clades within a larger clade of presumably free-living *Methanocorpusculum* species isolated from the environment, and are similar to them functionally as well (see below).

Overall, based on sequence identity and phylogeny, we recovered two major ciliate-associated clades, one exclusively *Plagiopyla-*associated and the other associated with both *Metopus* and *Plagiopyla*. Our previous marker gene-based analysis revealed that symbionts were specific at the host species level, but they did not form clades based on host lineage and their marker gene phylogeny is poorly resolved (19). We would expect these clades to be distinct considering the concurrent independent transitions to anoxia and symbiont acquisition events that are believed to have occurred across the phylum Ciliophora (52,53). It is possible that the close relationships observed between some *Plagiopyla*- and *Metopus*-associated strains are the consequence of symbiont switching/replacement events between these co-occurring hosts.

Notably, the environmental lineages which cluster with the ciliate symbionts originate from natural habitats, whereas the others were isolated from engineered systems (Figure S1). While classified as “environmental” here, it is important to note that most were obtained from mixed environmental samples and therefore may have originated from within protist cells. Only *M. bavaricum*, *M. labreanum Z*, and *M. parvum* XII represent cultured isolates, which are less likely to be symbiotic due to their ability to grow as free-living microbes. Still, since it is not yet known whether ciliate endosymbionts can grow outside of their hosts, we cannot completely exclude the possibility that these are also actually ciliate endosymbionts.

Altogether, our results demonstrate that *Methanocorpusculum* is composed of two divergent clades that originate from 1) animal-associated and 2) ciliate-associated or environmental habitats, spanning diverse environments including the digestive systems of rhinoceros, elephants, and ruminants, as well as the cytoplasm of anaerobic ciliates and a wide range of natural and engineered anoxic aquatic habitats. Although environmental *Methanocorpusculum* are most often reported from engineered systems, they have also been detected in marine habitats such as estuarine sediments, a stratified lagoon, hydrothermal vents, and particle-associated in the Cariaco Basin (54) (55–57). These occurrences may reflect undetected ciliate hosts rather than truly free-living populations, as has been hypothesized previously for *Methanoregula* identified in a freshwater lake (58). Given that diverse marine anaerobic ciliates harbor *Methanocorpusculum*, including the divergent taxa *Metopus*, *Plagiopyla*, and *Thigmothrix* (18) spanning three ciliate Classes, and the fact that this genus has only rarely been encountered in marine habitats which also harbor these ciliates, it is likely that *Methanocorpusculum* from marine habitats primarily exists as endosymbionts. Further environmental sampling and mining of existing datasets will be necessary to better understand the full habitat range and preferred lifestyle of this remarkably flexible genus.

### Animal- and ciliate-associated Methanocorpusculum species have distinct amino acid metabolic potential and host-related adaptations across disparate host habitats

We performed functional enrichment analysis to identify differences in metabolic potential across animal-, ciliate-associated, and environmental *Methanocorpusculum*. We observed significant differences in amino acid metabolism and nutrient acquisition, likely influenced by the disparate nature of and selective pressures within their various hosts. Prior analysis of animal-associated *Methanocorpusculum* also reported reduced biosynthetic capacity in animal-associated genomes relative to environmental lineages (16); our study extends this to ciliate endosymbionts, providing valuable insights into the impacts of divergent host habitats (i.e., living inside of a protist cell versus extracellularly within an animal gut) on symbiont genetic potential.

In our analysis, the ciliate-associated/environmental clade showed significantly broader amino acid biosynthetic potential than the animal-associated clade. We found that this clade is enriched in many more COG gene families and biosynthetic KEGG modules related to amino-acid metabolism and cofactor biosynthesis, many of which were entirely absent from the animal-associated genomes (Figure 3). These patterns highlight the pronounced biosynthetic potential of ciliate-associated and environmental *Methanocorpusculum* relative to the reduced metabolic repertoires of animal-associated lineages. In contrast, animal-associated *Methanocorpusculum* were enriched in several nutrient-salvage and catabolic gene families that were absent from the other groups. These include histidine transporter genes which may contribute to host-adapted nutrient acquisition that has been observed in other gut methanogens (59) and a di-/tri-peptidase gene which has previously been implicated in host-microbe interactions in the gut (60). Overall these results illustrate a shift in animal-associated *Methanocorpusculum* toward amino acid and nutrient salvaging in place of biosynthesis, likely due to supply of essential amino acids and cofactors present in host nutrition or from the metabolism of co-occurring gut microbes.

We also found other genomic differences that reflect divergent adaptations to host-associated environments. Ciliate-associated/environmental genomes were fully enriched in genes encoding proline/glycine transport enzymes (Figure 3) and glycan biosynthesis pathways (Figure 4). This potentially reflects management of osmotic stress tolerance and adaptation to the marine environment, as observed in *Methanocorpusculum labreanum* (61), as well as intracellular adhesion, as mannose-rich glycans and N-glycan rich structures are known to play roles in symbiont-host adhesion in microbial eukaryotes (19,62). In contrast, animal-associated genomes encoded functions related to bile-acid detoxification, a function that has been associated with gut-associated prokaryotes (63), and the 3’->5’ ssDNA/RNA exonuclease TatD, which plays a role in DNA repair under oxidative stress, likely important for survival in the animal gut (64).

In order to better understand the implications of these functions and their potential role in host adaptation, we compared *Methanocorpusculum* to related genera within Methanocorpusculaceae and Methanomicrobiales. These included the animal-associated *Methanorbis* and free-living *Methanocalculus* (17) (Figure S1), as well as *Methanofollis* and *Methanoculleus*, which are free-living and the most closely-related genera to Methanocorpusculaceae (17,22). *Methanocorpusculum* is particularly divergent from the rest of Methanomicrobiales, and so this allowed us to investigate changes in gene content during the evolutionary transition(s) to host association.

We found that many genomic features of animal-associated *Methanocorpusculum* were also shared with the insect, marsupial-, and chicken-gut associated, sister genus *Methanorbis*, including enrichment in transport and catabolic functions and the loss of biosynthetic pathways (e.g. tryptophan) (Figure S3). However, animal-associated *Methanocorpusculum* formed a distinct metabolic cluster (Figure S2), driven in part by further reductions in amino acid and cofactor biosynthesis, suggesting more extreme host specialization. Overall, these functional differences were unique to the animal-associated strains and were not present in the free-living relatives *Methanocalculus*, *Methanofollis*, and *Methanoculleus*, suggesting that they do indeed represent host adaptations. Notably, *Methanorbis* is hosted by termites and millipedes, which are known to harbor anaerobic protists with methanogenic symbionts (17,65). This raises the possibility that some *Methanorbis* lineages may be influenced by protist-rich gut niches, similar to the *Methanobrevibacter* endosymbiont of a termite-associated parabasalid, which has been suggested to undergo mixed-mode transmission in the gut environment (65). This may explain why, although they retain some animal-associated genomic traits discussed above, their genomes are functionally more similar to the protist-associated *Methanocorpusculum* than to the animal-associated clade.

Together, our findings demonstrate that *Methanocorpusculum* strains associated with animal or ciliate hosts exhibit genomic adaptations to host-association which vary dramatically by host type, even among animal hosts. Ciliate-associated strains retain broad biosynthetic and adhesive abilities, while the animal-associated strains appear to be streamlined for nutrient scavenging, stress tolerance, and potentially immune evasion.

### Ciliate-associated and environmental Methanocorpusculum share evolutionary trajectory and functional potential

We observed that ciliate-associated and environmental *Methanocorpusculum* strains are not only very similar phylogenetically (Figure 2a), but are also nearly indistinguishable functionally, sharing a nearly overlapping metabolic profile (Figure 2b). Very few COG functions and KEGG modules (Figures 4 and 5, Table S3) were enriched between these groups, and there were virtually no functions that were present in all ciliate-associated taxa that were absent in all environmental taxa, and vice versa.

This similarity, assuming that the environmental strains are truly free-living, is surprising considering the different selective pressures exerted on symbiotic versus free-living cells. It is thought that symbiont genomes undergo genome degradation and function loss when they are restricted to host tissue due to decreases in population size and homologous recombination, transmission bottlenecks and the resulting genetic drift, and fixation of deleterious mutations and loss of pseudogenes, ultimately leading to an organelle-like genome which is a fraction of the size of its free-living ancestors (66,67).

However, this pattern does not adequately describe many marine symbioses, in which symbiont gene flow via horizontal transmission and recombination between symbiotic lineages can maintain large genomes, even if the symbionts are vertically transmitted and rarely encounter genetically differentiated cells (66). Lind et al. 2018 (44) described a *Methanocorpusculum* symbiont of *Metopus* as being in an early stage of endosymbiont evolution, but it is possible that its mixed transmission mode prevents genome reduction by maintaining a reliable source of genomic variation via homologous recombination, and/or that dual adaptation to both host-associated and environmental habitats necessitates maintenance of these genes. Kaneko et al. (65) described a *Methanobrevibacter* endosymbiont of the parabasalid *Cononympha*, which lives within the hindgut of termites and might employ mixed-mode transmission between protist cells and the termite gut wall, potentially representing a strategy for response to changing conditions in the host gut. It is possible that the *Methanocorpusculum* symbionts of ciliates are similarly flexible, swapping between free-living and host-associated states in response to changing conditions within their host cells and the dynamic, intertidal sediments where they reside.

Indeed, a previous analysis indicated that *Methanocorpusculum* has likely experienced a higher rate of horizontal gene transfer (HGT) than other Methanomicrobiales genera and is the only genus in this Class to have experienced net gene loss. (61). This gene loss could potentially reflect the primarily host-associated lifestyle of *Methanocorpusculum*, as symbionts are expected to lose genes over time, while the high levels of HGT could be a result of mixed-mode transmission and/or dual adaptation to host-associated and environmental habitats and result in the maintenance of genomic diversity within symbiont populations. A larger analysis of gene loss and homologous recombination across Methanomicrobiales, with more host-associated and free-living lineages, is needed to investigate this further.

While high levels of homologous recombination with other symbiont and environmental lineages during mixed-mode transmission may explain the similarity that we observed between ciliate-associated and environmental *Methanocorpusculum*, there are other potential explanations and they are not mutually exclusive. First of all, as discussed above, it is possible that these free-living lineages are actually symbionts, or recently were symbionts, of ciliates or other anaerobic protists. Indeed, anaerobic ciliates and protists have been recovered in virtually all of the habitats that *Methanocorpusculum* has been found (Figure S1), including in wastewater (68,69), anaerobic digesters (70), groundwater (71), petroleum-polluted environments (72), estuarine sediment (19,20,73), and hydrothermal vents (74,75). In some of these habitats, such as hydrothermal vents, anaerobic ciliates are reported far more often than *Methanocorpusculum*, increasing the likelihood that *Methanocorpusculum* from this habitat are indeed endosymbiotic.

In our extended analysis, we recovered several potential signatures of host association which were present across all *Methanorbis* and *Methanocorpusculum*, including the *Methanocorpusculum* strains which are putatively environmental. For example, none of the genomes from these genera possessed the chemotaxis-related genes *cheW* and *cheY* or the archaeal flagellin gene *flaB* (Table S6) whereas *Methanocalculus*, *Methanofollis*, and *Methanoculleus* all did, consistent with a shared ancestral loss of motility and environmental sensing in the common ancestor of Methanocorpusculaceae (excluding *Methanocalculus*). On the other hand, some animal- and ciliate-associated *Methanocorpusculum* were missing biosynthetic cobalamin genes and KEGG modules which were present in the other lineages, including the environmental *Methanocorpusculum* (Figure S5). This represents the only major functional distinction between environmental and ciliate-associated *Methanocorpusculum* in our analysis. Loss of cobalamin biosynthesis may be expected in host-associated lineages due to the high energetic cost (76). Indeed, this absence has been observed in the genomes of the endosymbiotic methanogens *Methanobrevibacter* sp. NOE, *Methanocorpusculum* sp. MCE (44), and *Ca.* Methanobrevibacter cononymphae (65), suggesting that it may be a common trait of endosymbiotic methanogens. If the environmental populations are in fact endosymbionts of uncharacterized anaerobic protists, cobalamin biosynthesis may be selectively maintained by periodic exposure to cobalamin-limited conditions during horizontal transmission events or transient free-living phases between hosts.

Alternatively, assuming that the environmental lineages described here are indeed free-living, it is possible that their immediate environment is similar enough to the ciliate intracellular environment to produce the genomic similarities observed here. While the animal-associated *Methanocorpusculum* are housed deep within host tissues and physically separated from the external environment, the methanogenic endosymbionts of ciliates reside within ciliate cytoplasm and are exposed to the external environment after cell lysis and during occasional symbiont replacement events. Inside the ciliate cell, these symbionts are housed near host MROs, which provide them with the necessary substrates and an intracellular niche ideal for methanogenesis. The metabolic integration observed between methanogenic symbiont and anaerobic ciliate host is not dissimilar to the syntrophic relationships that methanogens form outside of a eukaryotic host (23,52,77). This symbiosis is beneficial to the symbionts particularly in marine environments, where they are normally outcompeted by sulfate-reducing bacteria for methanogenesis substrates (e.g. H_2_) (78). Here methanogens occupy a sulfate-limited niche that sulfate reducers cannot access; sulfate reducing bacteria can also be present as ectosymbionts where they have access to sulfate in marine water (52,77).

On the other hand, other prokaryotic symbionts of anaerobic ciliates have diverged considerably from their free-living counterparts. The denitrifying bacterial endosymbiont of a freshwater plagiopylean ciliate has a strongly reduced genome (79,80), and the same is true to some extent for the Firmicutes TC1 endosymbiont of *Trimyema compressum* (81) and the endosymbiont *Ca.* Hydrogeosomobacter endosymbioticus of anaerobic scuticociliate GW7 (82,83). This is also true for some bacterial endosymbionts of ciliates found in hypoxia: *Thiodyction*, the purple sulfur bacterial endosymbiont of *Psueodoblepharisma tenue*, is phylogenetically distinct from its free-living relatives and has a highly reduced genome and specialized metabolism with altered photosynthesis physiology (84). Therefore, it is likely that there is some feature of the methanogenic symbionts rather than the hosts that contributes to their similarity to environmental lineages.

Similar to *Methanocorpusculum*, the *Polynucleobacter* symbionts of the microaerophilic ciliate *Euplotes* are very closely-related to and encode genomes which are comparable in size with their free-living counterparts (85). They are thought to represent recently established symbionts undergoing the early stages of genome erosion which are ultimately destined for extinction and cyclical symbiont replacement (86,87). While there are fundamental differences in these symbiotic systems – there is no convincing evidence for this type of cyclical replacement in endosymbiotic *Methanocorpusculum* – the close relationships observed with the free-living relatives of both *Methanocorpusculum* and *Polynucleobacter* are striking. Free-living *Methanocorpusculum* genomes are small (1.7-1.8 Mb) (16) and streamlined, with a high degree of adaptation to a cooperative, syntrophic lifestyle necessary in energy-deficient anoxic habitats (23,88). Like *Methanocorpusculum*, free-living *Polynucleobacter* have small, streamlined genomes, which are specialized for life in oligotrophic environments (86). It is possible that the syntrophic nature of even free-living methanogens complicates the evolutionary trajectory of the symbionts and that they are pre-adapted to symbiosis with their already streamlined genomes (88), and similar may be true for *Polynucleobacter*. *Methanocorpusculum*, along with the other methanogenic endosymbionts of anaerobic protists, may be preadapted for host-associated mutualism due to their pre-existing integration into other microbial metabolisms.

## Conclusions

In this study, we conducted comprehensive pan- and phylogenomic analyses of 106 *Methanocorpusculum*, as well as 18 *Methanorbis*, 10 *Methanocalculus*, 15 *Methanoculleus*, and six *Methanofollis* genomes and MAGs, in order to investigate potential evolutionary and functional divergence across disparate host-associated and environmental habitats. Our findings provide insight into the host-associated adaptations of a prokaryotic genus which associates both extracellularly with animals and intracellularly with protists, a diversity of hosts which has rarely been observed outside of the methanogenic symbionts of protists and animals. We found that the animal-associated MAGs and the ciliate-associated and environmental clades formed two distinct, independently evolved clades and that all the *Methanocorpusculum* lineages studied here have some genomic adaptations to host-association which vary dramatically by host type. Animal-associated symbionts appear to be adapted to the gut environment, with enriched functions characterized by a reduction in amino acid and cofactor biosynthetic capabilities, nutrient scavenging strategies, and immune tolerance. Ciliate-associated and environmental *Methanocorpusculum* are virtually indistinguishable phylogenetically and functionally and retain broader biosynthetic functions, reflecting either more recent symbiont acquisition or the impact of mixed-mode transmission on maintaining stable, gene-rich symbioses without extensive genome reduction and gene loss. Our findings challenge the notion that intracellular lifestyles lead to genome erosion and gene loss and underscore the importance of transmission mode and host habitat in shaping symbiont evolution.

## Data Availability

Raw metagenomic data has been deposited to NCBI’s Sequence Read Archive under the BioProject PRJNA1457442. Symbiont MAGs have been submitted to GenBank under the accession numbers listed in Table S2.

## Acknowledgements

This work was supported by a Simons Foundation Early Career Investigator in Marine Microbial Ecology and Evolution Award to RAB, the Human Frontiers in Science Program, and funding from the Czech Grant Agency to IC (project no. 23-06004S).

## Author Contributions

**Anna Schrecengost** writing: original draft, review and editing; investigation, conceptualization, methodology, data curation, formal analysis, visualization. **Johana Rotterová:** writing: review and editing, methodology, investigation. **Ivan Čepička:** writing: review and editing, resources, supervision. **Roxanne A. Beinart:** funding acquisition, conceptualization, resources, supervision, writing: review and editing.

