## Supplemental Information and Figures for "Distinct genomic adaptations of the methanogenic archaeal genus *Methanocorpusculum* to symbiosis with animals and protists"

**Contents**

**EXPERIMENTAL PROCEDURES**

**REFERENCES**

**SUPLLEMENTAL FIGURES AND TABLES**

### SUPPLEMENTAL INFORMATION

### EXPERIMENTAL PROCEDURES

#### *Single ciliate cell isolation and DNA sequencing*

Single ciliate cells from *Metopus* populations and one *Plagiopyla* population (JUMA2P) were previously isolated and prepared for whole-genome shotgun sequencing (1). In this study, six *Plagiopyla* strains (Table S1) were isolated using the same protocol. Three to seven cells per strain were ultimately included in our analysis. Briefly, cells were picked with a glass micropipette under a stereomicroscope and washed eight times with sterilized seawater, or until no bacteria was visible in the wash medium. To obtain enough genetic material for shotgun sequencing, whole genome amplifications (WGA) were carried out on these single cells using the RepliG Advance DNA Single Cell kit (Qiagen) following the manufacturer's protocol. The resulting DNA was quantified with a Qubit fluorometer and submitted for Illumina NovaSeq6000 S4 150-bp paired-end shotgun sequencing SeqWell plexWell™ 96 NGS library kit.

#### *Bioinformatic assembly and binning of endosymbiont MAGs*

Illumina paired-end reads were quality trimmed and filtered using Trimmomatic (v0.39) (2) to remove adapter sequences and low-quality bases. These quality-filtered reads were aligned to a contaminant database using Bowtie2 (v2.5.2) (3) with default parameters, in order to remove contaminant reads. Reads which mapped to the contaminant database were excluded from downstream processes using samtools (v1.19.2) (4). Unmapped read IDs were extracted and used to recover clean reads from the original fastq files using seqtk (v1.4).

Paired-end reads were assembled *de novo* using metaSPAdes (v3.15.5) (5) with multiple k-mer values (21, 31, 41, 51, 61, 71, 82, 91) to maximize assembly quality. Coverage depth was calculated by mapping reads back to the assembled scaffolds with Bowtie2. The resulting alignment files were processed with samtools and METABAT2 (v2.15) (6) jgi_summarize_bam_contig_depths. Contigs were binned with both MetaBAT2 (v2.15; minimum contig length = 1500 bp) and MaxBin2 (v2.2.7; minimum contig length = 500 bp) (7). The resulting bins were refined using DAS Tool (v1.1.7) (8), integrating outputs from both binning approaches to produce consensus metagenome-assembled genomes (MAGs).

Symbiont bins were identified from our data via taxonomic assignment with GTDB-tk classify_wf (9): based on our previous knowledge of the symbionts of these ciliate populations (10), bins that  were assigned to the genus *Methanocorpusculum* were considered symbiont bins. Only bins with at least 80% completeness and less than 5% contamination as determined by checkM2 (11) were included in further analyses. All genomes and MAGs included in our analysis are described in Table S2 along with the associated quality statistics.

*Phylogenomic analyses of symbionts and phylogenetic analysis of hosts*

We inferred a maximum-likelihood phylogenomic tree based on the concatenated alignment of 220 single-copy core genes identified across the pangenome of 106 high-quality *Methanocorpusculum* MAGs and reference genomes. The tree was constructed using IQ-TREE with 1000 boostraps and the Whelan-And-Goldman evolutionary model. A maximum-likelihood phylogenomic tree was also generated in the same manner based on the concatenated alignment of 199 single-copy core genes identified across the pangenome of 85 high quality Methanomicrobiales MAGs and reference genomes.

We also constructed 18S rRNA phylogenetic trees of the host ciliate populations. 18S rRNA sequences of these populations were obtained previously (10,12), with the exception of *Plagiopyla* sp. SACTRIN, *Plagiopyla ovata* JUMA2P, and *Metopus* sp. JUMA2M, whose sequence was generated by performing BLASTn searches of its metagenomic assembly versus collections of *Plagiopyla* and *Metopus* 18S rRNA gene sequences (BLAST+ v2.15.0) (13). These sequences were aligned, along with reference sequences obtained from GenBank, using the MAFFT algorithm and the progressive methods G-INS-i. Alignments were manually trimmed to the primer regions using AliView (14) and phylogenetic trees were generated using a Maximum likelihood method in RaxML under the GTRGAMMAI model with 1000 bootstraps

### SUPPLEMENTAL TABLES

**Table S1**. Culture names and associated metadata.

**Table S2**. List of genomes included in pangenomic analysis of *Methanocorpusclum* and their associated quality statistics, as determined by CheckM2.

**Table S3**. Average nucleotide identity of *Methanocorpusclum* populations.

**Table S4**. Functional Enrichment Analysis of *Methanocorpusculum* pangenome.

**Table S5**. List of genomes included in extended pangenomic analysis of Methanocorpusculaceae and their associated quality statistics, as determined by CheckM2.

**Table S6.** Functional enrichment analysis of extended pangenomic analysis of Methanocorpusculaceae

### SUPPLEMENTAL FIGURES

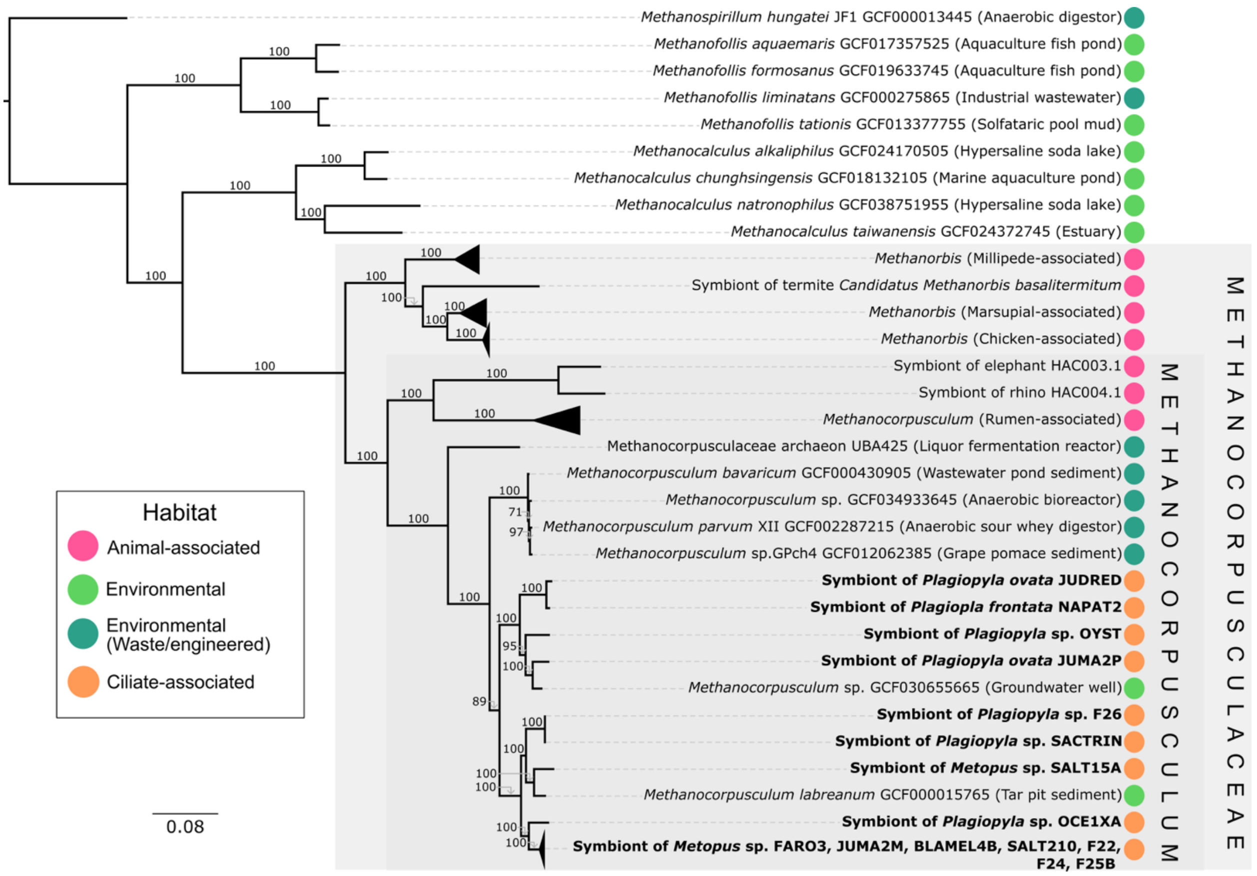


**Figure S1.** Phylogenomic tree of Methanocorpusculaceae, with symbionts of ciliates shown in bold. High quality genomes from NCBI RefSeq for the closely related Methanomicrobiales genera *Methanoculleus* and *Methanofollis* are also included as an outgroup. Nodes of the tree are

colored based on habitat, as pictured in the legend, and more detailed descriptions of habitats are included in the node labels. The maximum-likelihood phylogenomic tree was inferred from a concatenated alignment of 199 single-copy core genes (SCCGs) identified across 85 Methanomicrobiales genomes and constructed using 1000 bootstraps (only bootstrap values above 80 are shown on branches). Scale bar indicates substitutions per site.

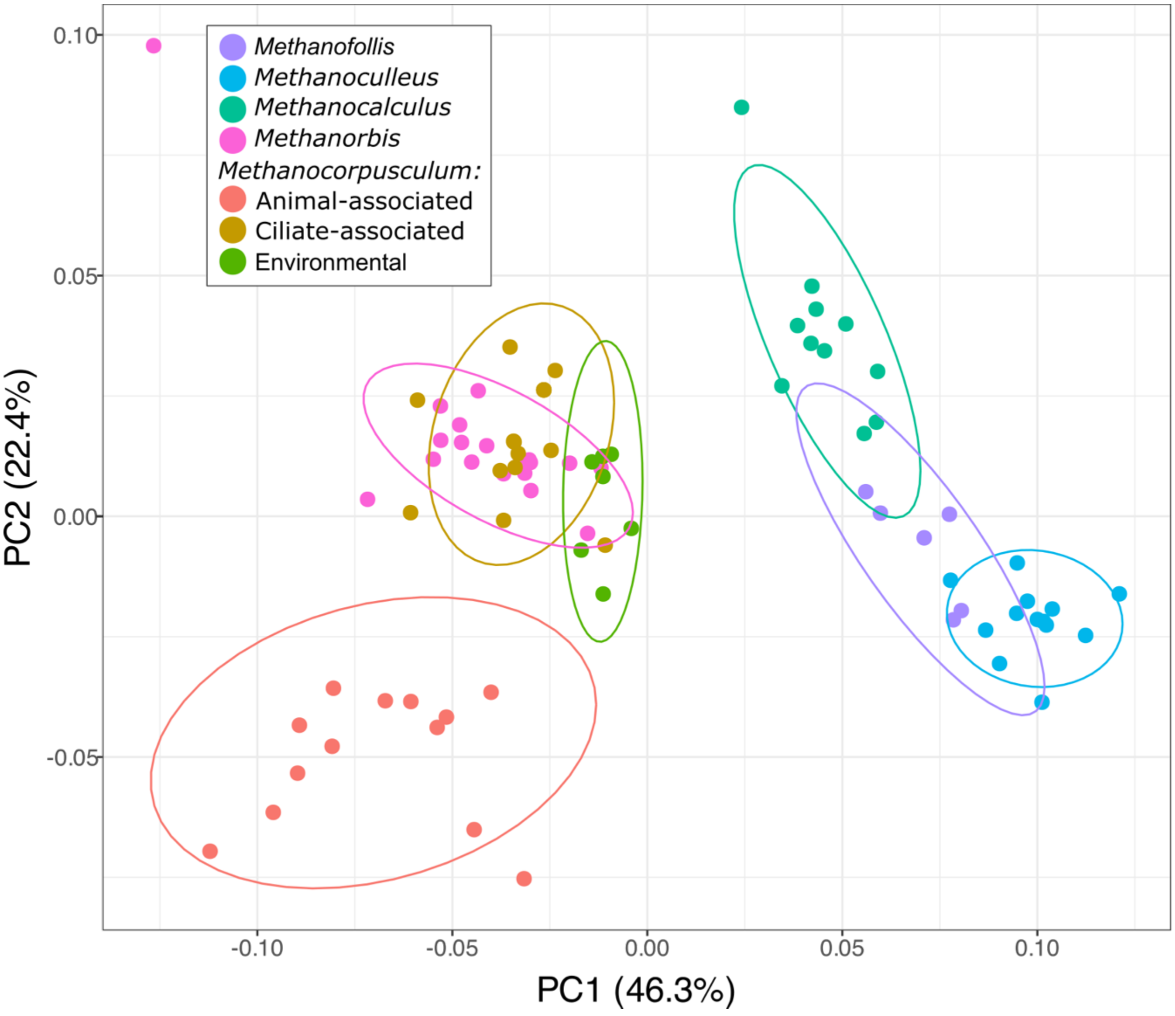


**Figure S2.** PCoA analysis based on Bray-Curtis distances calculated from pathway completeness profiles of KEGG modules in the supplemental pangenomes. Points represent genomes or MAGs and *Methanocorpusculum* genomes/MAGs are colored by habitat while the others are colored by genus. Axes show percentage of variance explained: PC1: 46.3%, PC2: 22.4%. Ellipses represent 95% confidence interval for grouping by habitat/genus.

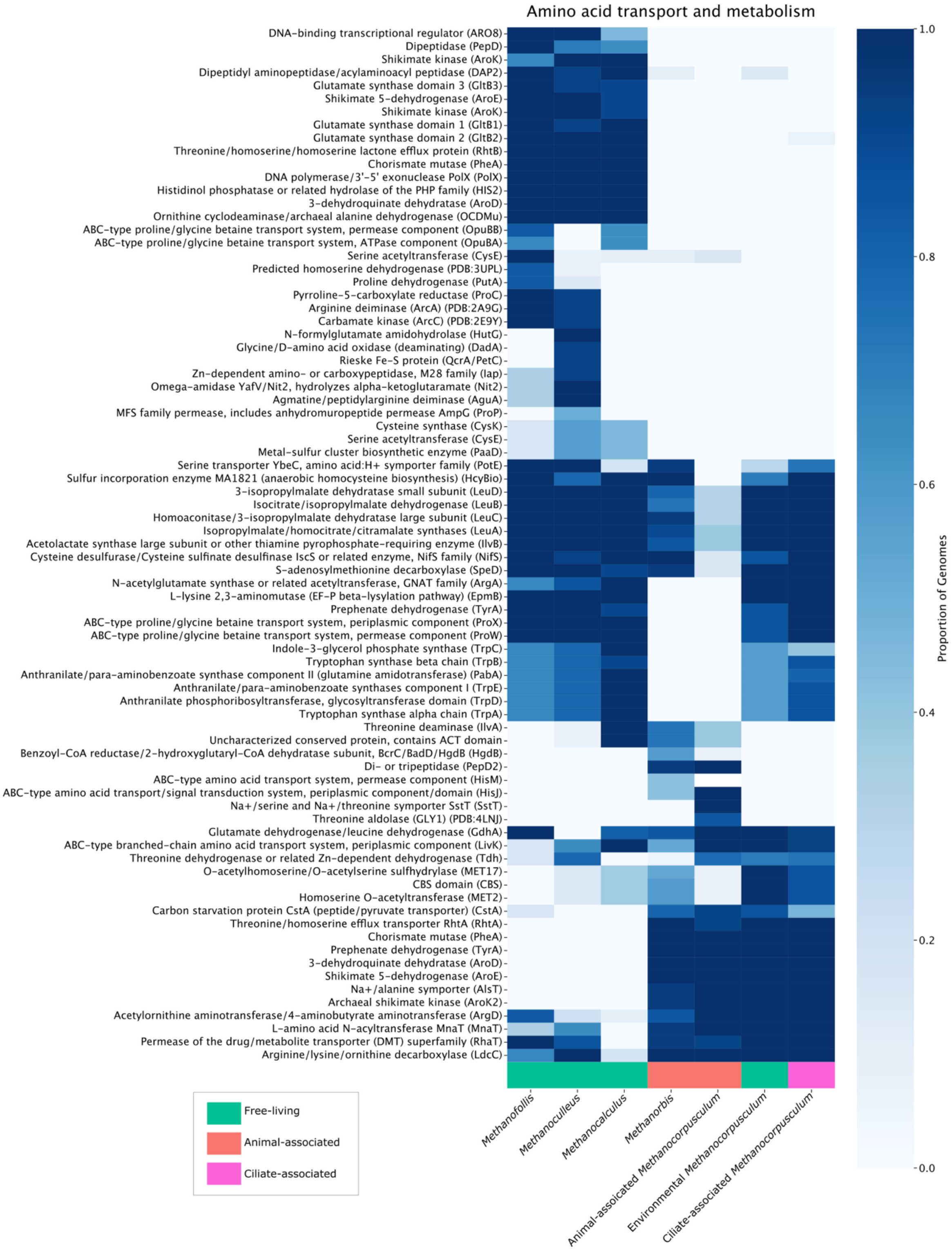

**Figure S3**. Functionally enriched enzymes involved in amino acid metabolism across the supplemental pangenome. *Methanocorpusculum* genomes/MAGs were grouped by habitat and the rest grouped by genus. These groups are colored at the bottom of the figure based on habitat: free-living, animal-, or ciliate-associated. Gene clusters were annotated with COG functions. Color indicates the proportion of genomes in each group which contain a given COG function. Only enzymes with a functional enrichment score of at least 30 were included, and all of these are significantly enriched ($p<0.05$).

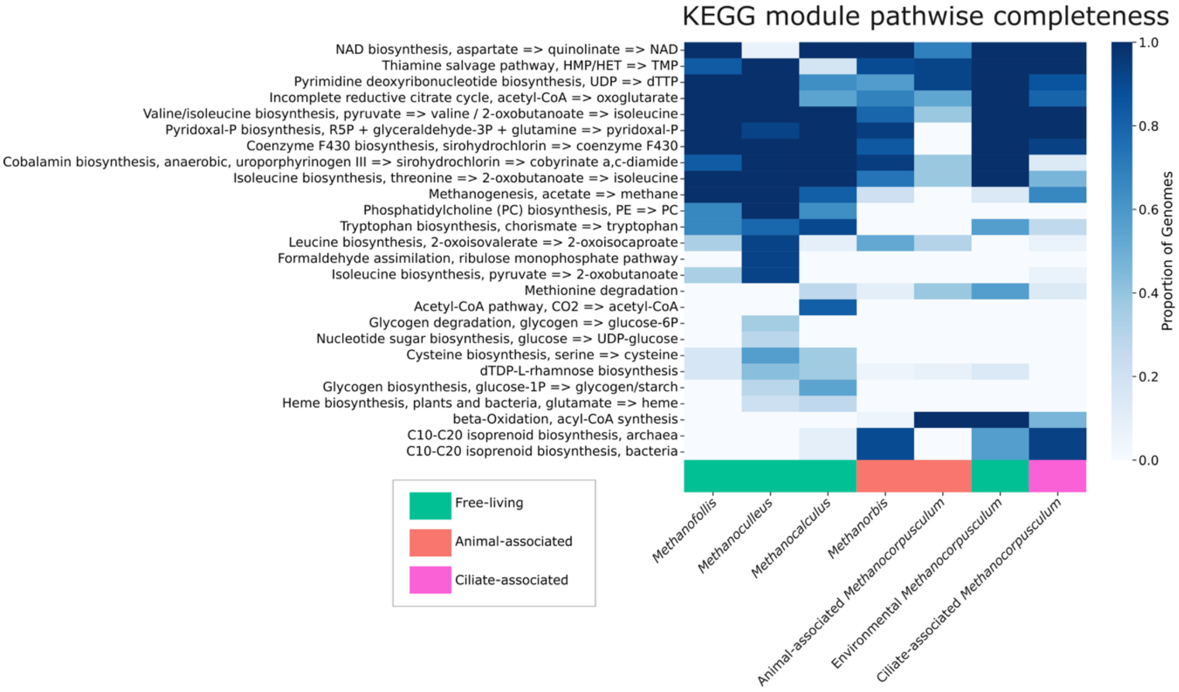


**Figure S4**. Functionally enriched KEGG modules based on their pathwise completeness scores across the supplemental pangenome. *Methanocorpusculum* genomes/MAGs were grouped by habitat and the rest grouped by genus. These groups are colored at the bottom of the figure based on habitat: free-living, animal-, or ciliate-associated. Gene clusters were annotated with COG functions. Color indicates the proportion of genomes in each group which contain a given COG function. Only modules with a functional enrichment score of at least 30 were included, and all of these are significantly enriched ($p<0.05$).

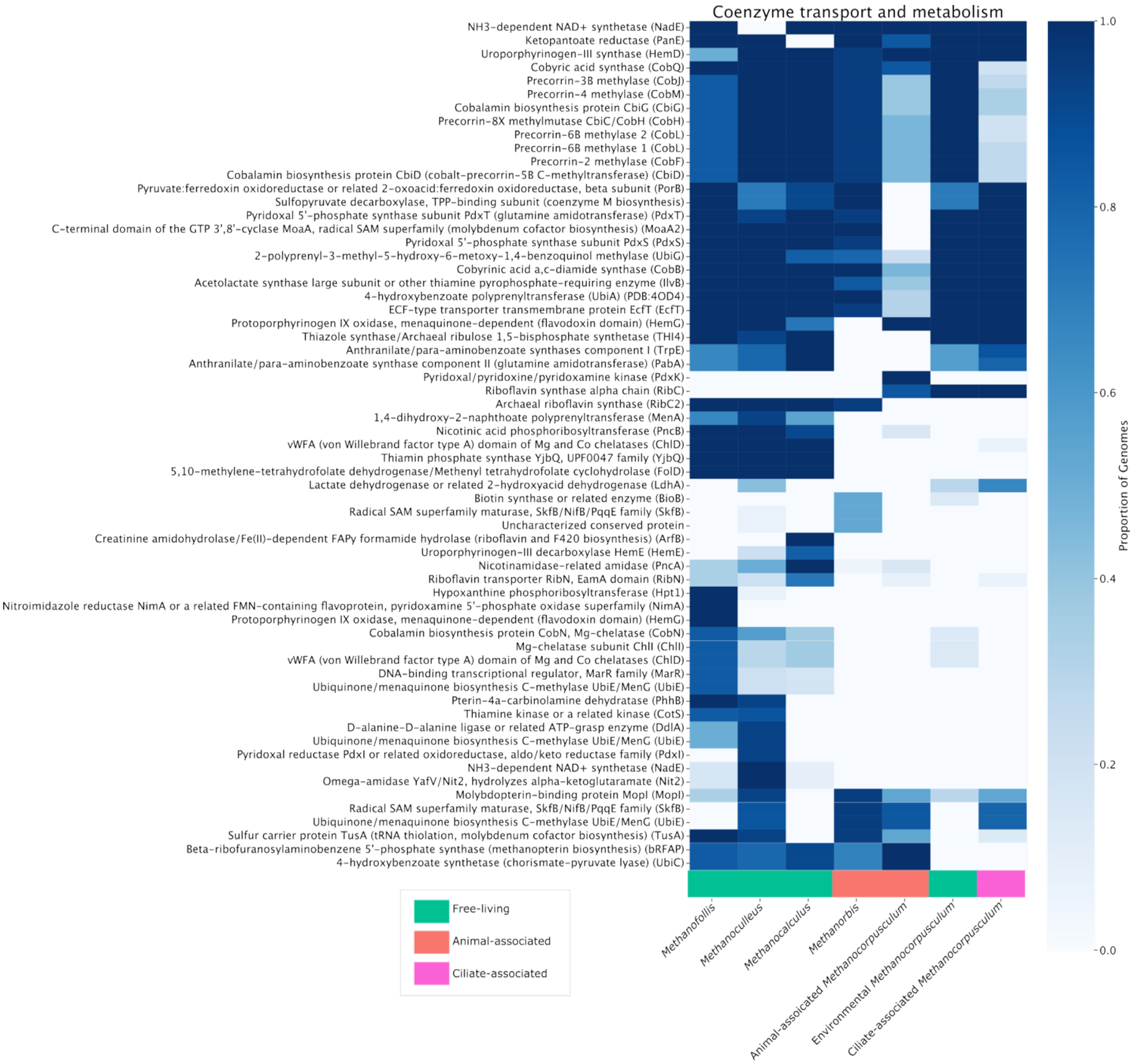


**Figure S5**. Functionally enriched enzymes involved in cofactor transport and metabolism across the supplemental pangenome. *Methanocorpusculum* genomes/MAGs were grouped by habitat and the rest grouped by genus. These groups are colored at the bottom of the figure based on habitat: free-living, animal-, or ciliate-associated. Gene clusters were annotated with COG functions. Color indicates the proportion of genomes in each group which contain a given COG function. Only enzymes with a functional enrichment score of at least 30 were included, and all of these are significantly enriched ($p$<0.05).
